# NEON Crowns: a remote sensing derived dataset of 100 million individual tree crowns

**DOI:** 10.1101/2020.09.08.287839

**Authors:** Ben. G. Weinstein, Sergio Marconi, Stephanie Bohlman, Alina Zare, Aditya Singh, Sarah J. Graves, Ethan White

## Abstract

Forests provide essential biodiversity, ecosystem and economic services. Information on individual trees is important for understanding the state of forest ecosystems but obtaining individual-level data at broad scales is challenging due to the costs and logistics of data collection. While advances in remote sensing techniques allow surveys of individual trees at unprecedented extents, there remain significant technical and computational challenges in turning sensor data into tangible information. Using deep learning methods, we produced an open-source dataset of individual-level crown estimates for 100 million trees at 37 sites across the United States surveyed by the National Ecological Observatory Network’s Airborne Observation Platform. Each canopy tree crown is represented by a rectangular bounding box and includes information on the height, crown area, and spatial location of the tree. Tree crowns identified using this technique correspond well with hand-labeled crowns, exhibiting both high levels of overlap and good correspondence in height estimates. These data have the potential to drive significant expansion of individual-level research on trees by facilitating both regional analyses at scales of ~10,000 ha and cross-region comparisons encompassing forest types from most of the United States.

## Introduction

Trees are central organisms in maintaining the ecological function, biodiversity and the health of the planet. There are estimated to be over three trillion individual trees on earth (Crowther et al., 2015) covering a broad range of environments and geography (Hansen et al., 2013). Counting and measuring trees is central to developing an understanding of key environmental and economic issues and has implications for global climate, land management and wood production. Field-based surveys of trees are generally conducted at local scales (~0.1-100 ha) with measurements of attributes for individual trees within plots collected manually. Connecting these local scale measurements at the plot level to broad scale patterns is challenging because of spatial heterogeneity in forests. Many of the key processes in forests, including change in forest structure and function in response to disturbances such as hurricanes and pest outbreaks, and human modification through forest management and fire, occur at scales beyond those feasible for direct field measurement.

Satellite data with continuous global coverage have been used to quantify important patterns in forest ecology and management such as global tree cover dynamics and disturbances in temperate forests (e.g. Bastin et al., 2018). However, the spatial resolution of satellite data makes it difficult to detect and monitor individual trees that underlie large scale patterns. These shortcomings can however be overcome by utilizing higher resolution remotely sensed data from low Earth orbit satellites, aircraft or drones to capture individual-level changes in forest structure and composition (Aubry-Kientz et al., 2019; Puliti et al., 2020). These high-resolution data have become increasingly accessible but converting the data into information on individual trees requires significant technical expertise and access to high-performance computing environments. This prevents most ecologists, foresters, and managers from engaging with large scale data on individual trees, despite the availability of the underlying data products and broad importance for forest ecology and management.

In response to the growing need for publicly available and standardized airborne remote sensing data over forested ecosystems, the National Ecological Observatory Network (NEON) is collecting multi-sensor data for more than 40 sites across the US. In this research, we combine these sensor data with a semi-supervised deep learning approach (Weinstein et al., 2020b, 2019) to produce a dataset on the location, height and crown area of over 100 million individual canopy trees at 37 sites distributed across the United States. To make these data readily accessible, we are releasing easy to access data files to spur biological analyses and to facilitate model development for tasks that rely on individual tree prediction. We describe the components of this open-source dataset, compare predicted crowns with hand-labeled crowns for a range of forest types, and discuss how this dataset can be used to address key questions in forest research.

## The NEON Crowns dataset

The NEON Crowns dataset contains tree crowns for all canopy trees (those visible from airborne remote sensing) at 37 NEON sites. Since subcanopy trees are not visible from above, they are not included in this dataset. We operationally define “trees” as plants over 3m tall. The 37 NEON sites represent all NEON sites containing trees with co-registered RGB and LiDAR data from 2018 or 2019 (see S3 for a list of sites and their locations). Predictions were made using the most recent year for which images were available for each site.

The dataset includes a total of 104,675,304 million crowns. Each predicted crown includes data on the spatial position of the crown bounding box, the area of the bounding box (an approximation of crown area), the 99th quantile of the height of LiDAR returns within the bounding box above ground level (an estimate of tree height), the year of sampling, the site where the tree is located, and a confidence score indicating the model confidence that the box represents a tree. The confidence score can vary from 0-1, but based on results from (Weinstein et al., 2020b), boxes with less than 0.15 confidence were not included in the dataset.

The dataset is provided in two formats: 1) as 11,000 individual files each covering a single 1km^2 tile (geospatial ‘shapefiles’ in standard ESRI™ format); and 2) as 37 csv files, each covering an entire NEON site. Geospatial tiles have embedded spatial projection information and can be read in commonly available GIS software (e.g., ArcGIS, QGIS) and geospatial packages for most common programming languages used in data analysis (e.g., R, Python). All data are publicly available, openly licensed (CC-BY), and permanently archived on Zenodo (https://zenodo.org/deposit/3765872).

To support the visualization of the dataset have developed a web visualization tool using the ViSUS WebViewer (www.visus.org) to allow users to view all of the trees at the full site scale with the ability to zoom and pan to examine individual groups of trees down to a scale of 20m (see http://visualize.idtrees.org, Figure 2). This tool will allow the ecological community to assist in identifying areas in need of further refinement within large area covered by the 37 sites.

## Crown Delineation Methods

The location of individual tree crowns was estimated using a semi-supervised deep learning workflow (Figure 3; Weinstein et al., 2020b, 2019). This workflow uses a one-shot object detector with a convolutional neural network backbone to identify trees in RGB imagery. The model was pre-trained using weak labels generated from a previous published LiDAR tree detection algorithm using NEON data from 30 sites (Silva et al., 2016). The model was then trained on 10,000 hand-annotated crowns from 7 NEON sites (Figure 1). This phase of the workflow was performed using the DeepForest python package (Weinstein et al., 2020a). We extend the workflow by filtering trees using the LiDAR-derived canopy height model to remove objects identified by the model with heights of <3m (Supplementary Material). This addition was important in sparsely vegetated landscapes, such as oak savannah and deserts where it was difficult for the model to distinguish between trees and low shrubs in the RGB imagery.

**Figure 1.**
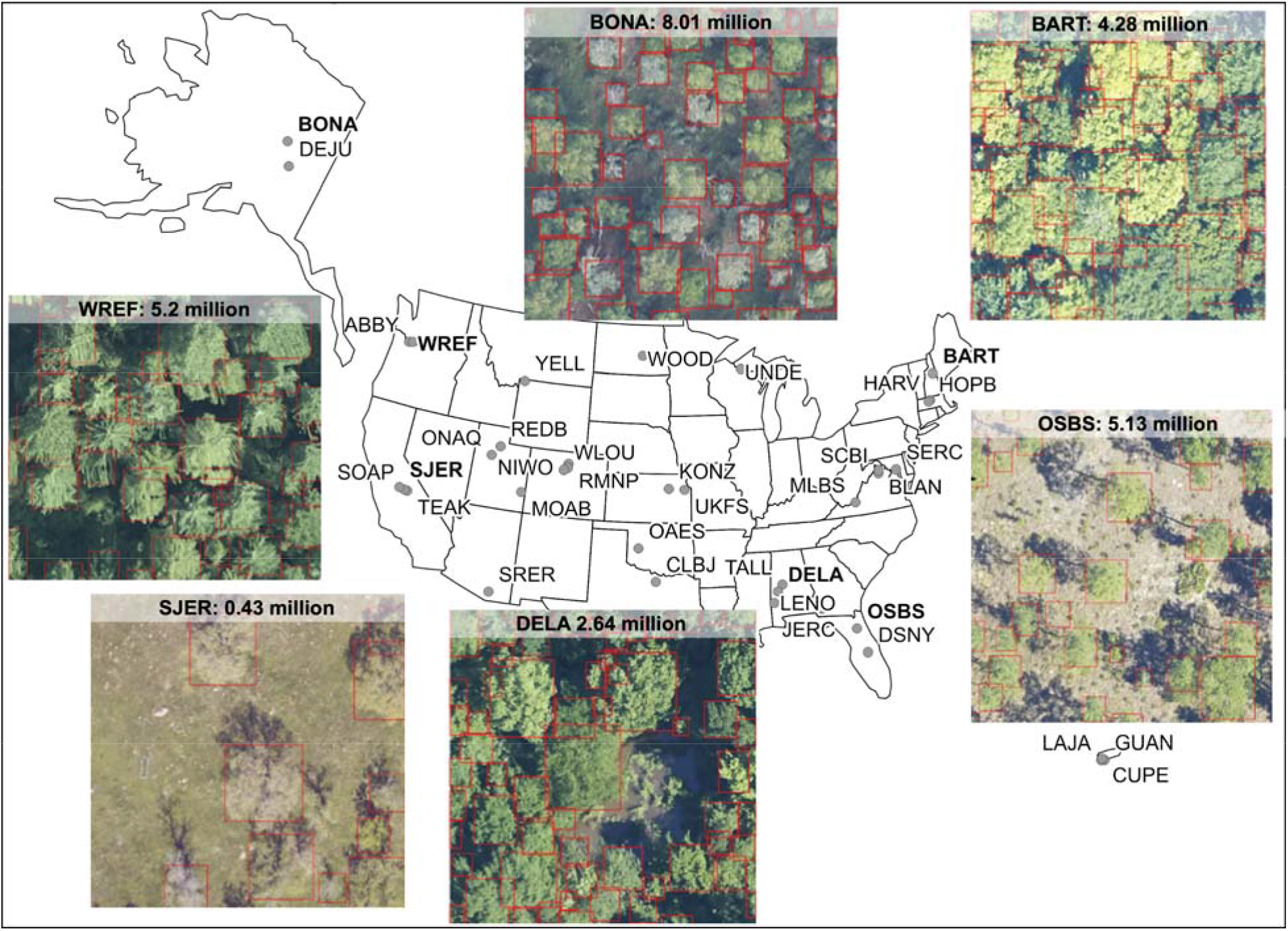
Locations of 37 NEON sites included in the NEON Crowns Dataset and examples of tree predictions shown with RGB imagery for six sites. Clockwise from bottom right: 1) OSBS: Ordway-Swisher Biological Station, Florida 2) DELA: Dead Lake, Alabama, 3) SJER: San Joaquin Experimental Range, California, 4) WREF: Wind River Experimental Forest, Washington, 5) BONA: Caribou Creek, Alaska and 6) BART: Bartlett Experimental Forest, New Hampshire. Each predicted crown is associated with the spatial position, crown area, maximum height estimate from co-registered LiDAR data, and a predicted confidence score.

**Figure 2.**
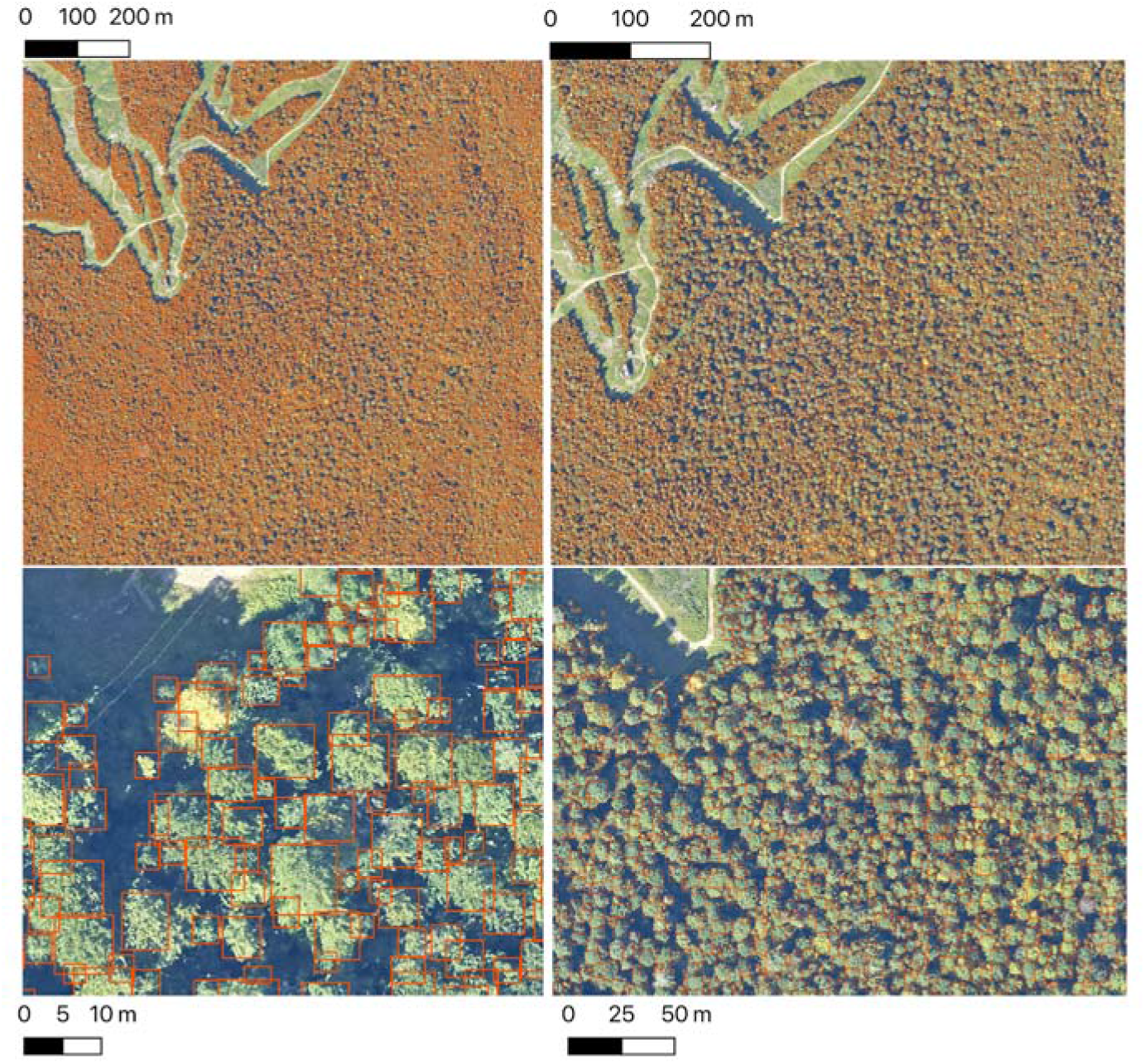
The Neon Crowns Dataset provides individual-level tree predictions at broad scales. An example from Bartlett Forest, NH shows the ability to continuously zoom from landscape level to stand level views. A single 1km tile is shown. NEON sites tend to have between 100 to 400 tiles in the full airborne footprint.

**Figure 3.**
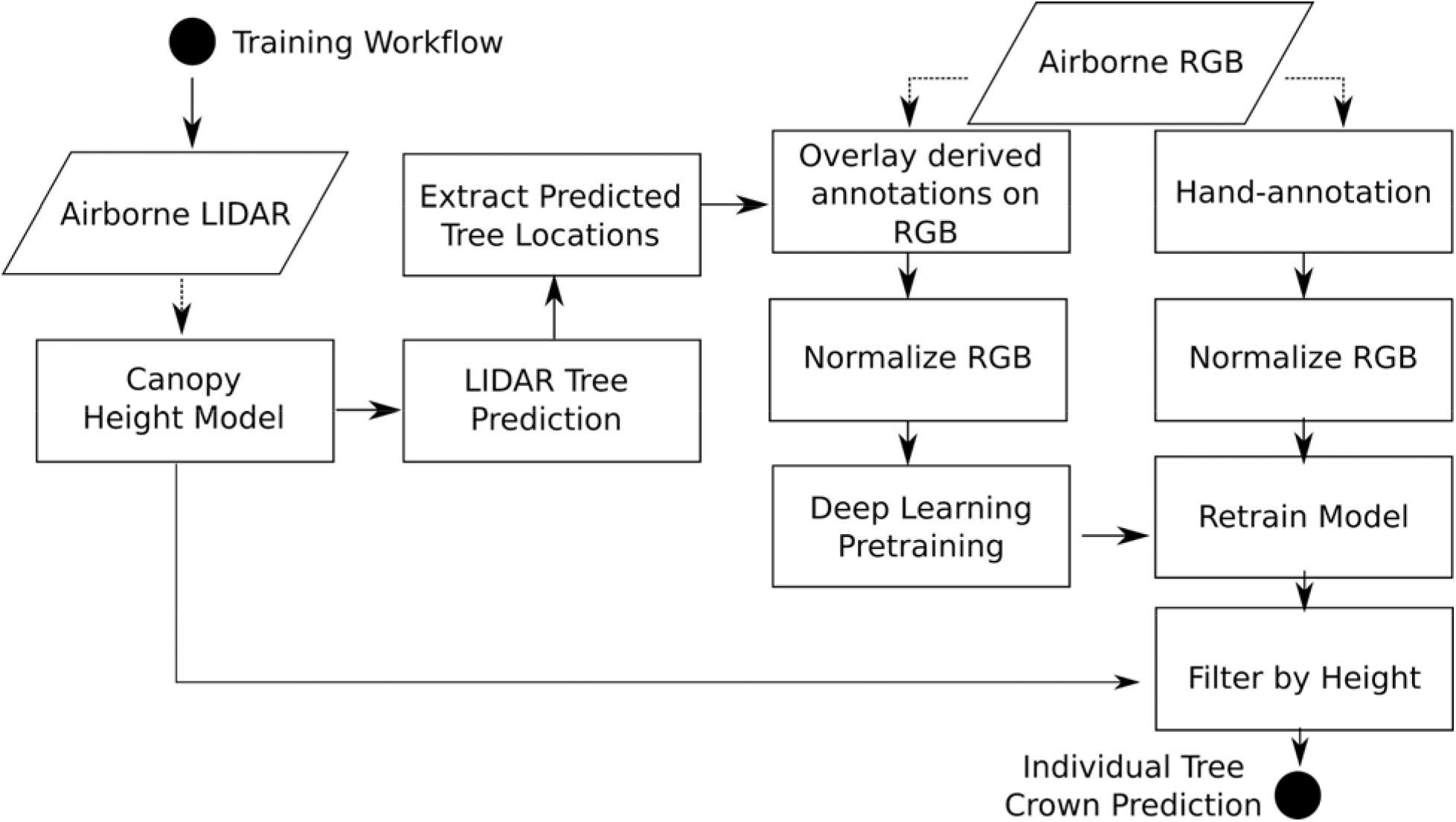
Workflow diagram adapted from (Weinstein et al., 2020a). The workflow for model training and development are identical to (Weinstein et al., 2020a) with the exception of extracting heights from the canopy height model for each bounding box prediction.

## Evaluation and Validation

Building on evaluation methods from Weinstein et al., (2020b, 2020a, 2019), we validated the dataset using hand-annotated bounding boxes drawn by an observer looking directly at the sensor data. We refer to this type of evaluation data as ‘image-annotated crowns’. This approach allows the performance of the crown-delineation algorithm to be evaluated across the full range of forest types represented in the continental-scale dataset. However, note that these image-annotated crowns will not be as accurate as field-annotated crowns (S. Graves et al., 2018), where an observer records crown position while physically next to the target tree. Image-annotated crowns may therefore overestimate the performance of the algorithm relative to more precise ground truth.

We compared predicted tree crowns to image-annotated crowns from 21 NEON sites (n=207 images, 6926 trees) that were withheld from model training. These sites were selected to cover a wide range of forest types and geographies. Using a 50% intersection over union threshold, our workflow yielded a bounding box recall of 72.4% with a precision of 70.5%. Recall is the proportion of image-annotated crowns matched to a crown prediction and precision is the proportion of predictions that match image-annotated crowns. Precision and recall are equally important for developing a tree crown dataset, because it is important to both successfully identify trees and ignore non-tree objects. Tests indicate that the model generalizes well across geographic sites and forest conditions (Figure 4; Weinstein et al., 2020a, 2020b), but there is a general bias towards undersegmenting trees in dense stands where multiple individual trees with similar optical characteristics are grouped into a single delineation. Additional training data and the LiDAR threshold added in this implementation resulted in predictions that were 4.1% more precise, but 2.8% less accurate than (Weinstein et al., 2020a) (Figure 4). The decrease in recall likely occurs because the NEON field plots that were used for evaluation occur mostly in forested areas of the NEON sites, rather than in less dense areas of the sites. Areas with less dense forest (e.g., agriculture, suburban areas, and bare ground) are not as common within the NEON field plots used for evaluation and are likely the areas with improved precision from the use of the new LiDAR threshold (Supplementary Material). The 4% increase in precision is therefore likely a lower bound and is worth the trade-off in the minimal drop in recall.

**Figure 4.**
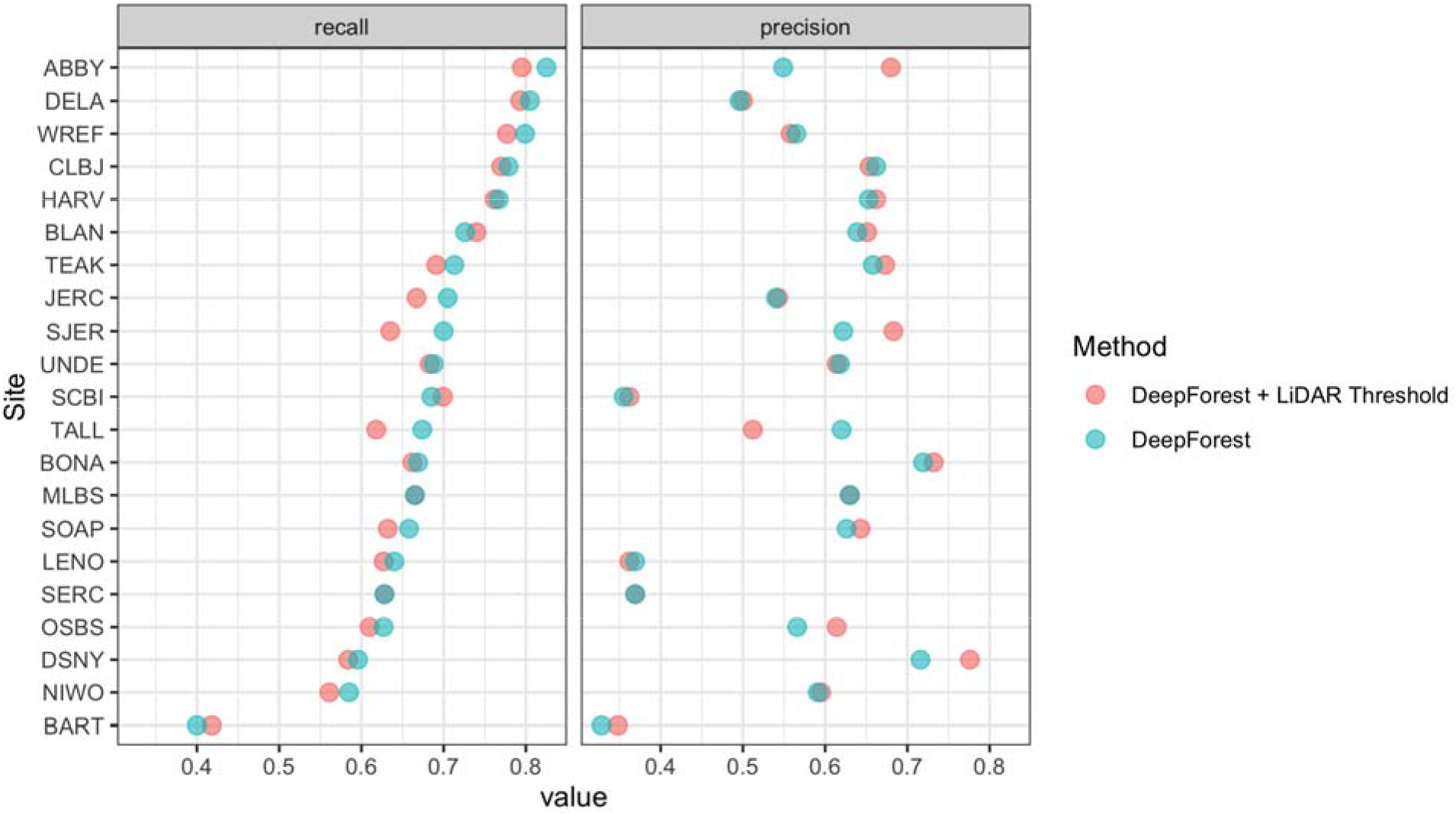
Precision and recall scores for the algorithm used to create the NEON Crowns Dataset (red points), as well as the DeepForest model from Weinstein et al., (2020a) (blue points). Evaluation is performed on 207 image-annotated images (6926 trees) in the NEONTreeEvaluation dataset (https://github.com/weecology/NeonTreeEvaluation).

We also compared crowns delineated by the algorithm to field-collected stems from NEON’s Woody Vegetation Structure dataset. This data product contains a single point for each tree with a stem diameter ≥ 10cm. We filtered the raw data to only include trees likely to be visible in the canopy (see Appendix S1). These overstory tree field data help us analyze the performance of our workflow in matching crown predictions to individual trees by scoring the proportion of field stems that fall within a prediction. Field stems can only be applied to one prediction, so if two predictions overlap over a field stem, only one is considered a positive match. This test produces an overall stem recall rate at 69.4%, which is similar to the bounding box recall rate from the image-annotated data (Figure 5). The analysis of stem recall rate is conservative due to the challenge of aligning the field-collected spatial data with the remote sensing data (Appendix S1). We found several examples of good predictions that were counted as false positives due to errors in the position of the ground samples within the imagery.

**Figure 5.**
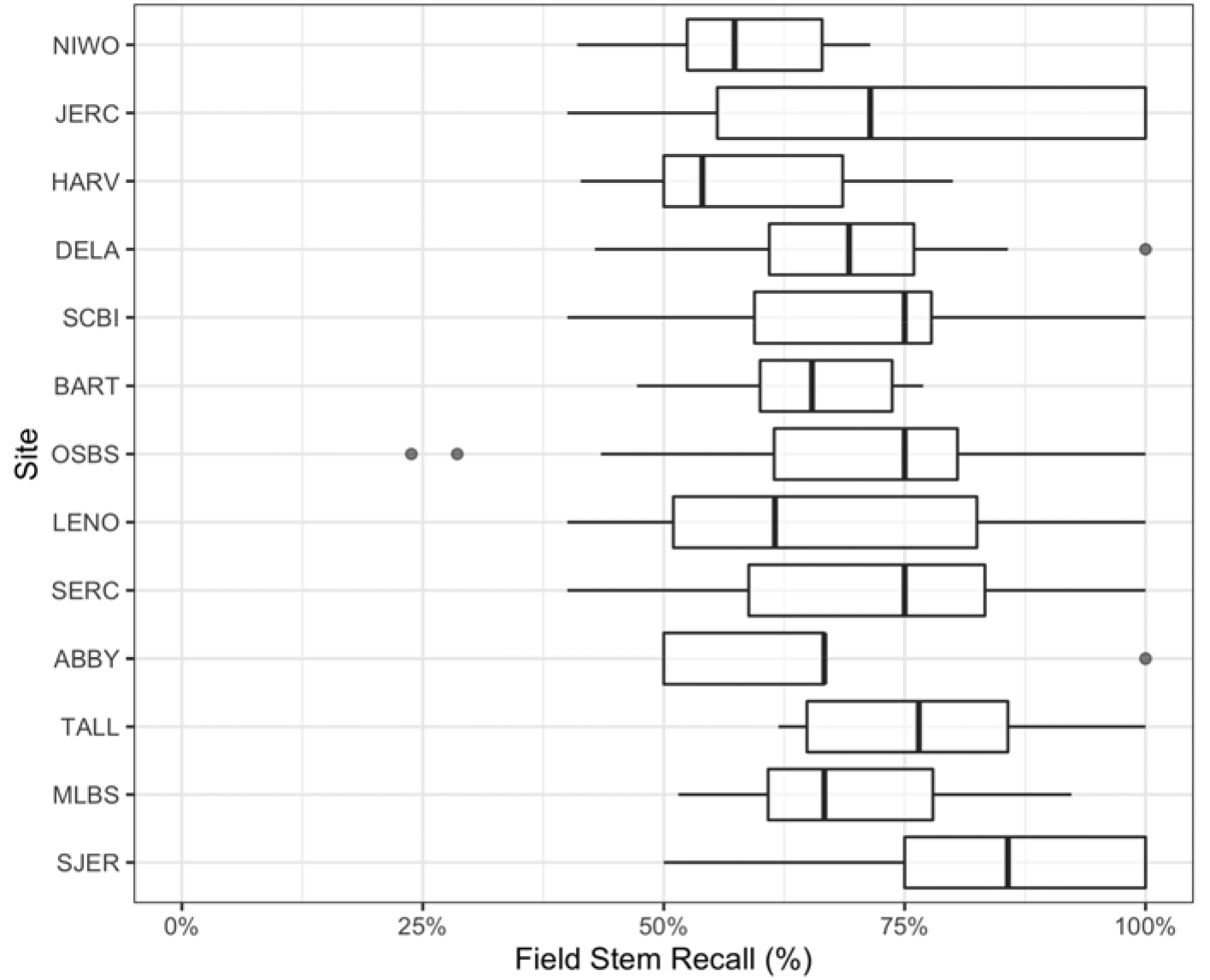
Overstory stem recall rate for NEON sites with available field data. Each data point is the recall rate for a field-collected plot. NEON plots are either 40mx40m ‘tower’ plots with two 20×20m subplots, or a single 20mx20m ‘distributed’ plot. See NEON sampling protocols for details. For site abbreviations see S3.

To assess the utility of our approach for mapping forest structure, we compared remotely sensed predictions of maximum tree height to field measurements of tree height of overstory trees using NEON’s Woody Plant Vegetation Structure Data. We used the same workflow described in Appendix S1 for determining overstory status for both the stem recall and height verification analysis. Predicted heights showed good correspondence with field-measured heights of reference trees. Using a linear-mixed model with a site-level random effect, the predicted crown height had a Root Mean Squared Error of 1.73m (Figure 6). The relationship is stronger in forests with more open canopies (SJER, OSBS) and predictably more prone to error in forests with denser canopies (BART, MLBS). Given the challenges of measuring tree heights, including the difficulty of measuring tree height in the field, the potential for tree growth between the time of field measurement and image acquisition (often several years), and the automated workflow to designate whether field-collected trees were visible in the canopy, these results suggest that overstory height measures are reasonably accurate across the dataset.

**Figure 6.**
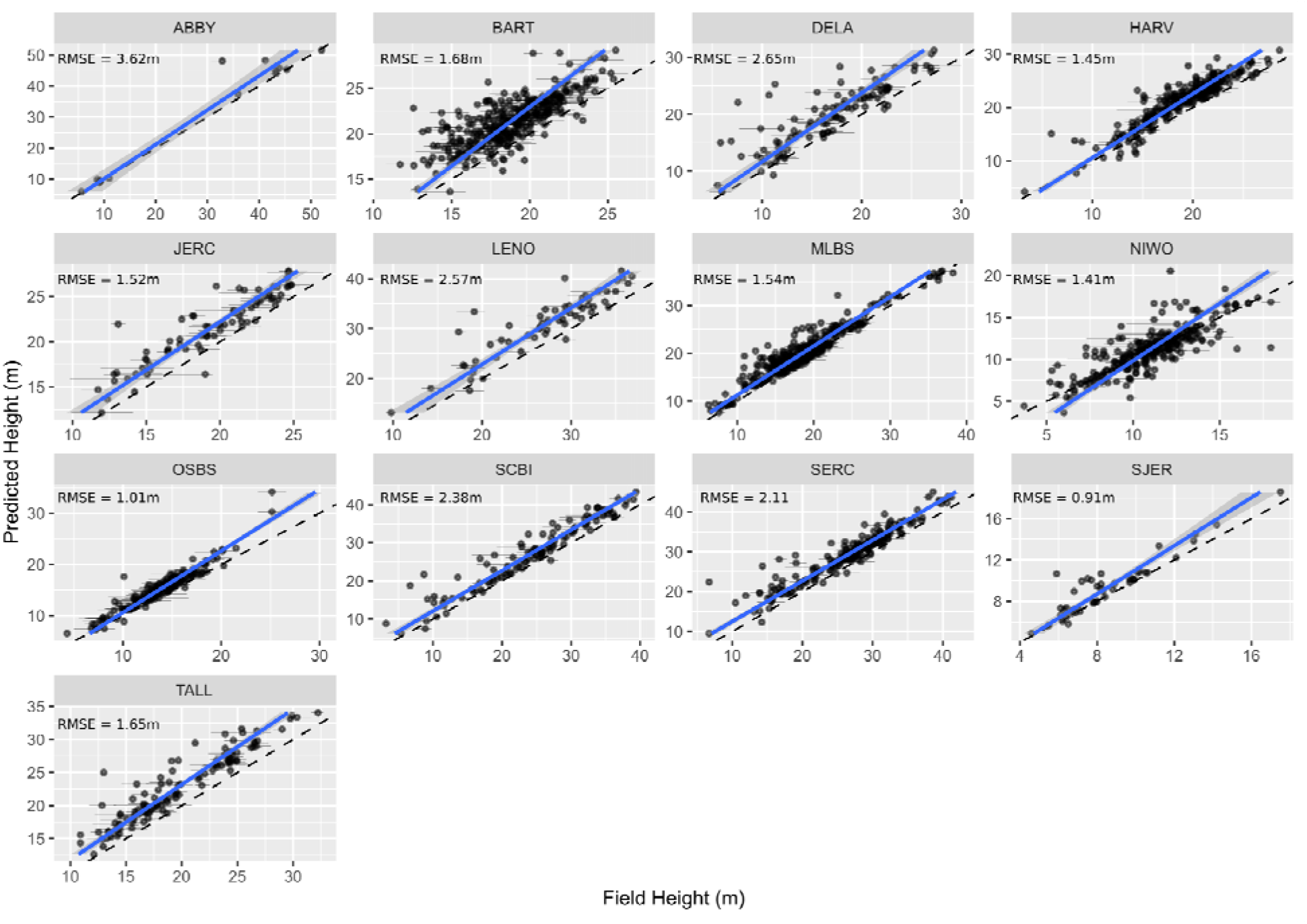
Comparison of field and remote sensing measurements of tree heights for 11 sites in the National Ecological Observatory Network. Each point is an individual tree. See text and S1 for selection criteria and matching scheme for the field data. The RMSE of a mixed-effects model with a site level random effect is 1.73m.

## Using the NEON Crowns dataset for individual, landscape and biogeographic scale applications

This dataset supports individual-level cross-scale ecological research that has not been previously possible. It provides the unique combination of information spanning the entire United States, with sites ranging from Puerto Rico to Alaska, with continuous individual-level data within sites at scales hundreds of times larger than what is possible using field-based survey methods. At the individual level, high-resolution airborne imagery can inform analysis of critical forest properties, such as tree growth and mortality (Clark et al., 2004), foliar biochemistry (Chadwick and Asner, 2016), and landscape-scale carbon storage (S. J. Graves et al., 2018). Because field data on these properties are measured on individual trees, individual level tree detection allows connecting field data directly to image data. In addition, growth, mortality and changes in carbon storage occur on the scale of individual trees such that detection of individual crowns allows direct tracking of these properties across space and time. While it is possible to aggregate information at the stand level, we believe that individual level data opens new possibilities in large scale forest monitoring and provides richer insights into the underlying mechanisms that drive these patterns.

At landscape scales, research is often focused on the effect of environmental and anthropogenic factors on forest structure and biodiversity. For example, understanding why tree abundance and biomass vary across landscapes has direct applications to numerous ecological questions and economic implications (e.g. Laubhann et al., 2009). Often this requires sampling at a number of disparate locations and either extrapolation to continuous patterns at landscape scales, or assumptions that the range of possible states of the system are captured by the samples. Using the individual level data from this dataset, we can now produce continuous high-resolution maps across entire NEON sites for enabling landscape scale studies of multiple ecological phenomena (Figure 7). These landscape scale responses can then be combined with high resolution data on natural and anthropogenic drivers (e.g., topography, soils, fire management) to model forest dynamics at broad scales.

**Figure 7.**
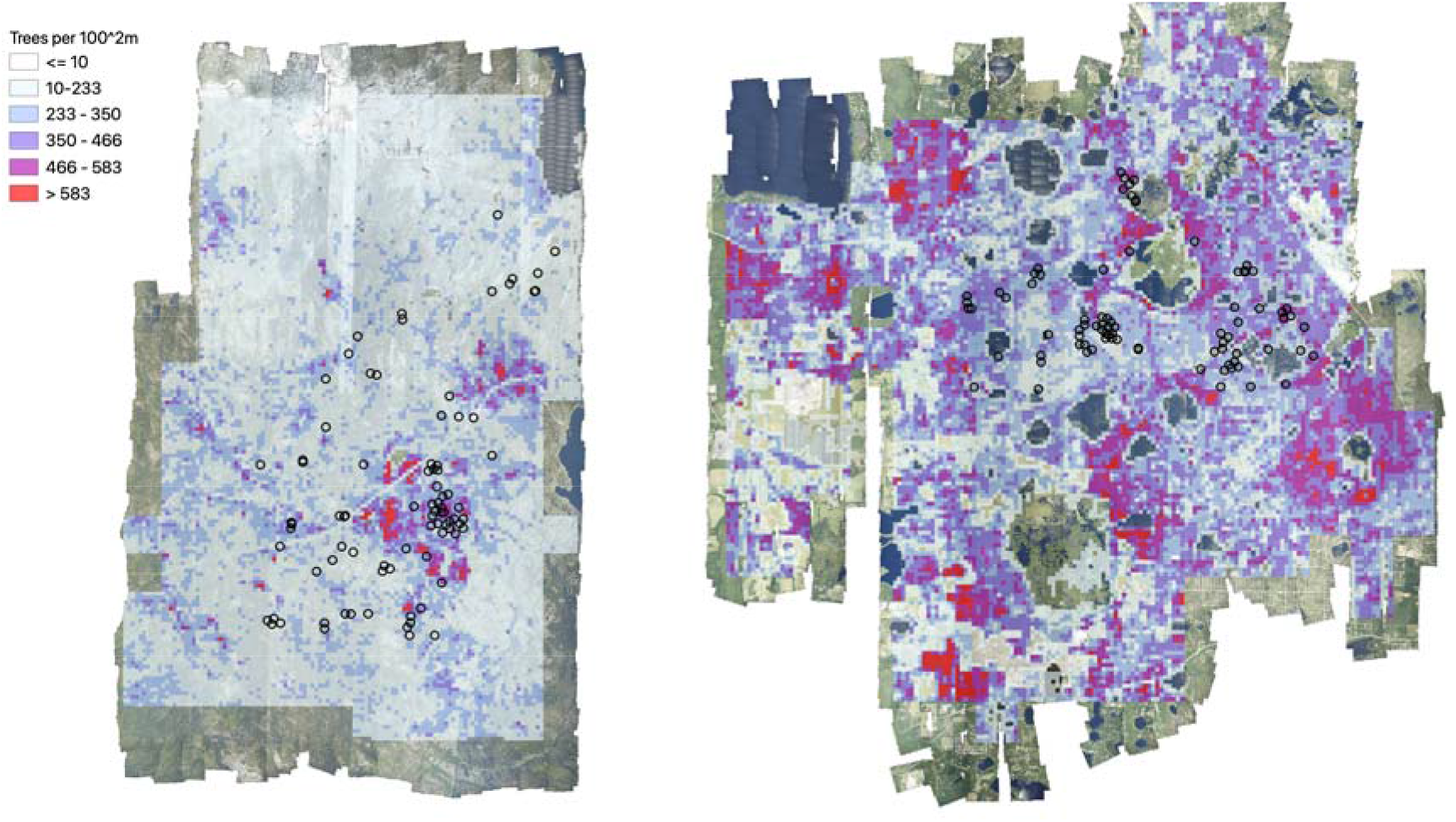
Tree density maps for Teakettle Canyon, California (left) and Ordway Swisher Biological Station, Florida (right). For each 100m^2 pixel, the total number of predicted crowns were counted. The location of NEON Woody Plant Vegetation sampling plots are shown in black circles.

By focusing on detecting individual trees, this approach to landscape scale forest analysis does not require assumptions about spatial similarity, sufficiently extensive sampling, or consistent responses of the ecosystem to drivers across spatial gradients. This is important because the heterogeneity of forest landscapes makes it difficult to use field plot data on quantities such as tree density and biomass to extrapolate inference to broad scales (Marvin et al., 2014). To illustrate this challenge, we compared field-measured tree densities of NEON field plots to estimated densities of 10,000 remotely sensed plots of the same size placed randomly throughout the landscapes across footprints of the airborne data. We attempted to change the Woody Vegetation data as little as possible (i.e. compared to the more refined filtered data in previous analyses) in order to obtain estimates of tree cover in a plot from the field data. To be included in this analysis, trees needed to have valid spatial coordinates and a minimum height of 3m. Some older data lacked height estimates, in which case we used a minimum dbh threshold of 15cm. In each simulated plot, we then counted the total number of predicted tree crowns to create a distribution of tree densities at the site level (Figure 8). Comparing the field plot tree densities with the distribution from the full site shows deviations for most sites, with NEON field plots exhibiting higher tree densities than encountered on average in the airborne data for some sites (e.g.,Teakettle Canyon, CA) and lower tree densities than from remote sensing in others (e.g., Ordway-Swisher Biological Station). While this kind of comparison is inherently difficult due to differing thresholds and filters for data inclusion in field versus remotely sensed data, it highlights that even well stratified sampling of large landscapes as was done with NEON plots (see NEON technical documents for NEON.DP1.10098) can produce differing tree attribute estimates than continuous sampling from remote sensing data. Combining representative field sampling with remote sensing to produce data products like the NEON Crowns dataset provides an approach to addressing this challenge to improve estimations of the abundance, biomass, and size distributions across large geographic areas.

**Figure 8.**
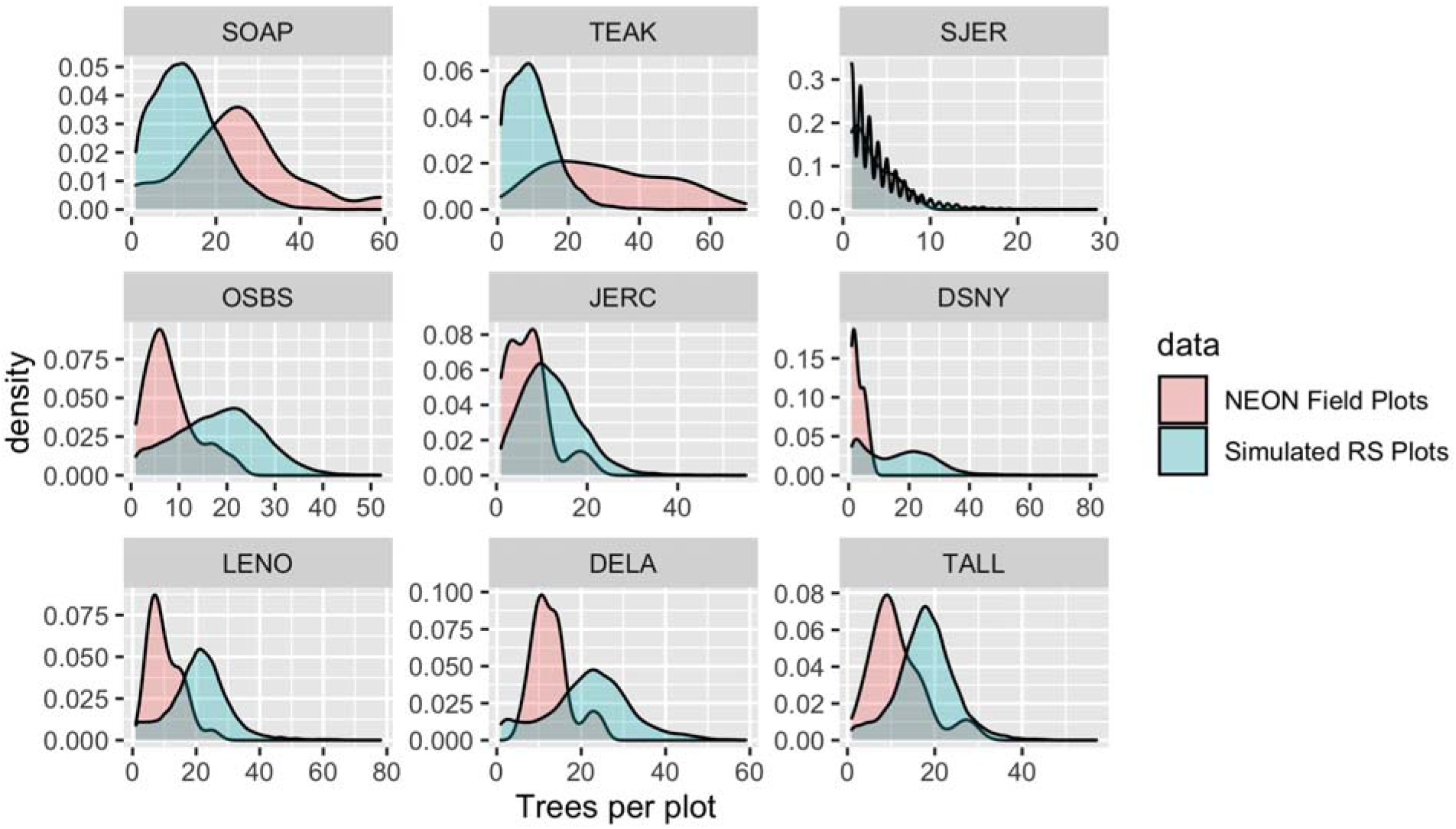
Comparison of tree counts between the field-collected NEON plots and the predicted plots from the dataset. For the remote sensing data, 10000 simulated 40m^2^ plots were calculated for each site for the full AOP footprint for each year. To mimic NEON sampling, 2 quadrants were randomly sampled in each simulated plot. No plots on water, bare ground, or herbaceous land classes were included in the comparison. We selected three sites from three NEON domains to show a sample of sites across the continental US. Both distributed and tower NEON plots were used for these analyses.

The NEON Crowns dataset supports the assessment of cross-site patterns to help understand the influence of large-scale processes on forest structure at biogeographic scales. For example, ecologists are interested in how and why forest characteristics such as abundance, biomass, and allometric relationships vary among forest types (e.g. Jucker et al., 2017) and how these patterns covary across environmental gradients (Feldpausch et al., 2011). Understanding these relationships is important for inferring controls over forest stand structure, understanding individual tree biology, and assessing stand productivity. By providing standardized data that span near-continental scales, this dataset can help inform the fundamental mechanisms that shape forest structure and dynamics. For example, we can calculate tree allometries (e.g., the ratio of tree height to crown area) on a large number of individual trees across NEON sites and explore how allometry varies with stand density and vegetation type (Figure 9). This example analysis shows a continental-scale relationship, with denser forests exhibiting trees with narrower crowns for the same tree height compared to less dense forests, but also clustering and variation in the relationship within vegetation types. For example, subalpine forests illustrate relationships between tree density and allometry that are distinct from other forest types. By defining both general biogeographic patterns, and deviations therein, this dataset will allow the investigation of factors shaping forest structure at macroecological scales.

**Figure 9.**
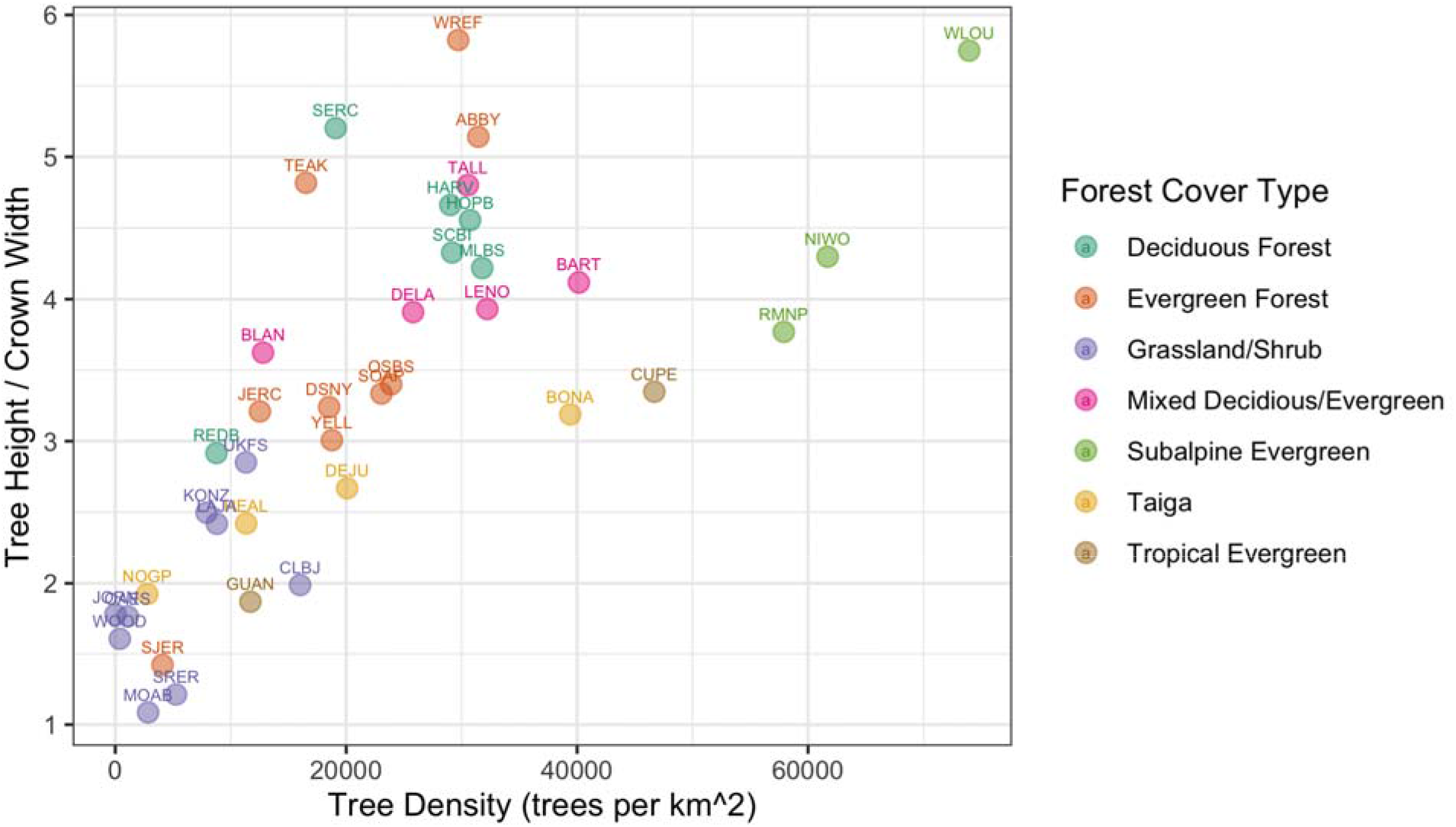
Individual crown attributes for predictions made at each NEON site. For site abbreviations see S1. Crown area is calculated by multiplying the width and height of the predicted crown bounding box. Crown height is the 99th quantile of the LiDAR returns that fall inside the predicted crown bounding box. Sites are colored by the dominant forest type to illustrate the general macroecological relationship among sites in similar biomes.

In addition to these ecological applications, the NEON Crowns dataset can also act as a foundation for other machine learning and computer vision applications in forest informatics, such as tree health assessments, species classification, or foliar trait estimation. In each of these tasks, individual tree delineation is the first step to associate sensor data with ground measurements. However, delineation requires a distinct set of technical background and computational approaches and thus many ecological applications either skip an explicit delineation step entirely (Williams et al., 2020) or apply existing software without detailed exploration of segmentation performance (e.g. Maschler et al., 2018). Ignoring these factors can hamper accurate assessments due to mismatches between sensor data and individuals. While our crown annotations are not perfect, they are specifically tailored to one of the largest and openly accessible datasets that allows pairing individual tree detections with information on species identity, tree health, and leaf traits through NEONs field sampling, and we believe they are sufficiently robust to serve as the basis for broad scale analysis.

## Limitations and Further Technical Developments

An important limitation for this dataset is that it only provides information on sun-exposed tree crowns. It is therefore not appropriate for ecological analyses that depend on accurate characterization of subcanopy trees and the three-dimensional structure of forest stands. Fortunately, a number of the major questions and applications in ecology are primarily influenced by large individuals (Enquist et al., 2020). For example, biomass estimation is largely driven by the canopy in most ecosystems, rather than mid or understory trees that are likely to be missed by aerial surveys. Similarly, habitat classification and species abundance curves can depend on the dominant forest structure that can be inferred from coarse resolution airborne data (Shirley et al., 2013) and could be improved using this dataset. It may be possible to establish relationships between understory and canopy measures using field data that could allow this dataset to be used as part of a broader analysis (Bohlman, 2015). However, this would require significant additional research to validate the potential for this type of approach.

An additional limitation is the uncertainty inherent in the algorithmic detection of crowns. While we found good correspondence between image-based crown annotations and those produced by the model for many sites, there remained substantial uncertainty across all sites and reasonable levels of error in some sites. It is important to consider how this uncertainty will influence the inference from research using this and similar datasets. The model is biased towards undersegmentation, meaning that multiple trees are prone to being grouped as a single crown. It is also somewhat conservative in estimating crown extent wherein it tends to ignore small extensions of branches from the main crown. These biases could impact studies of tree allometry and biomass if the analysis is particularly sensitive to crown area. When making predictions for ecosystem features such as biomass, it will be important to propagate the uncertainty in individual crowns into downstream analyses. While confidence scores for individual detections are provided to aid uncertainty propagation, the use of additional ground truth data may also be necessary to infer reliability.

One aspect of individual crown uncertainty that we have not addressed is the uncertainty related to image-based crown annotations and measurement of trees in the field (S. Graves et al., 2018). To allow training and evaluating the model across a broad range of forest types, we used image-based crown annotations. This approach assumes that crowns identifiable in remotely sensed imagery accurately reflect trees on the ground. This will not always be the case, as what appears to be a single crown from above may constitute multiple neighboring trees, and conversely, what appears to be two distinct crowns in an image may be two branches of a single large tree (S. Graves et al., 2018). Targeted field surveys will be always needed to validate these predictions and community annotation efforts will allow for assessment of this component of uncertainty.

The machine learning workflow used to generate this dataset also has several areas that could be improved for greater accuracy, transferability and robustness. The current model contains a single class ‘Tree’ with an associated confidence score. This representation prevents the model from differentiating between objects that are not trees and objects for which sufficient training information is not available. For example, the model has been trained to ignore buildings and other vertical structures that may look like trees. However, when confronted by objects data that has never been encountered, it often produces unintuitive results. Examples of objects that did not appear in the training data, and as a result were erroneously predicted as trees, include weather stations, floating buoys, and oil wells. Designing models that can identify outliers, anomalies, and ‘unknown’ objects is an active area of research in machine learning and will be useful in increasing accuracy in novel environments. In addition, NEON data can sometimes be afflicted by imaging artifacts due to co-registration issues with LiDAR and raw RGB imagery (Appendix S2). This effect can lead to distorted imagery that appears fuzzy and swirled and lead to poor segmentation. An ideal model would detect these areas of poor quality and label them as ‘unknown’ rather than attempting to detect trees in these regions.

Given these limitations, we view this version of the dataset as the first step in an iterative process to improve cross-scale individual level data on trees. Ongoing assessment of these predictions using both our visualization tool and field-based surveys will be crucial to continually identify areas for improvements in both training data and modeling approaches. While iterative improvements are important, the accuracy of the current predictions illustrates that this dataset is sufficiently precise for addressing many cross-scale questions related to forest structure. By providing broad scale crown data we hope to highlight the promising integration between deep learning, remote sensing, and forest informatics, and provide access to the results of this next key step in ecological research to the broad range of stakeholders who can benefit from these data.

## Acknowledgements

We would like to thank NEON staff and in particular Tristan Goulden and Courtney Meier for their assistance and support. This research was supported by the Gordon and Betty Moore Foundation’s Data-Driven Discovery Initiative (GBMF4563) to E.P. White and by the National Science Foundation (1926542) to E.P. White, S.A. Bohlman, A. Zare, D.Z. Wang, and A. Singh. The funders had no role in study design, data collection and analysis, decision to publish, or preparation of the manuscript.

