## Supplementary material for "NEON Crowns: a remote sensing derived dataset of 100 million individual tree crowns": S1-S5

### Supplemental Materials

### S1. Evaluation of the workflow on NEON collected field stems

Raw NEON tree stem data was processed from the NEON data portal to provide a data set to compare image derived heights to field measured heights. The following filters were applied to the raw NEON field data (ID: DP1.10098.001) after download. A reference tree must have

- Valid spatial coordinates
- A unique height measurement per sampling period. Individuals double recorded but with different heights were discarded
- Height measurements in more than one year to verify height measurement
- Changes in between-year field heights of less than 6m
- A classification as alive
- A minimum height of 3m to match the threshold in the remote sensing workflow.
- Be at least within 5m of the canopy as measured by the LiDAR height model extracted at the stem location. This was used to prevent matching with understory trees in the event that overstory trees were eliminated due to failing in one of the above conditions, or not sampled by NEON.

To match trees in the field and the NEON Crowns dataset, we took the closest height when two predictions and field stems overlapped. We also dropped CLBJ since only 3 points met this criteria. All other NEON sites did not have any data that met this criteria.

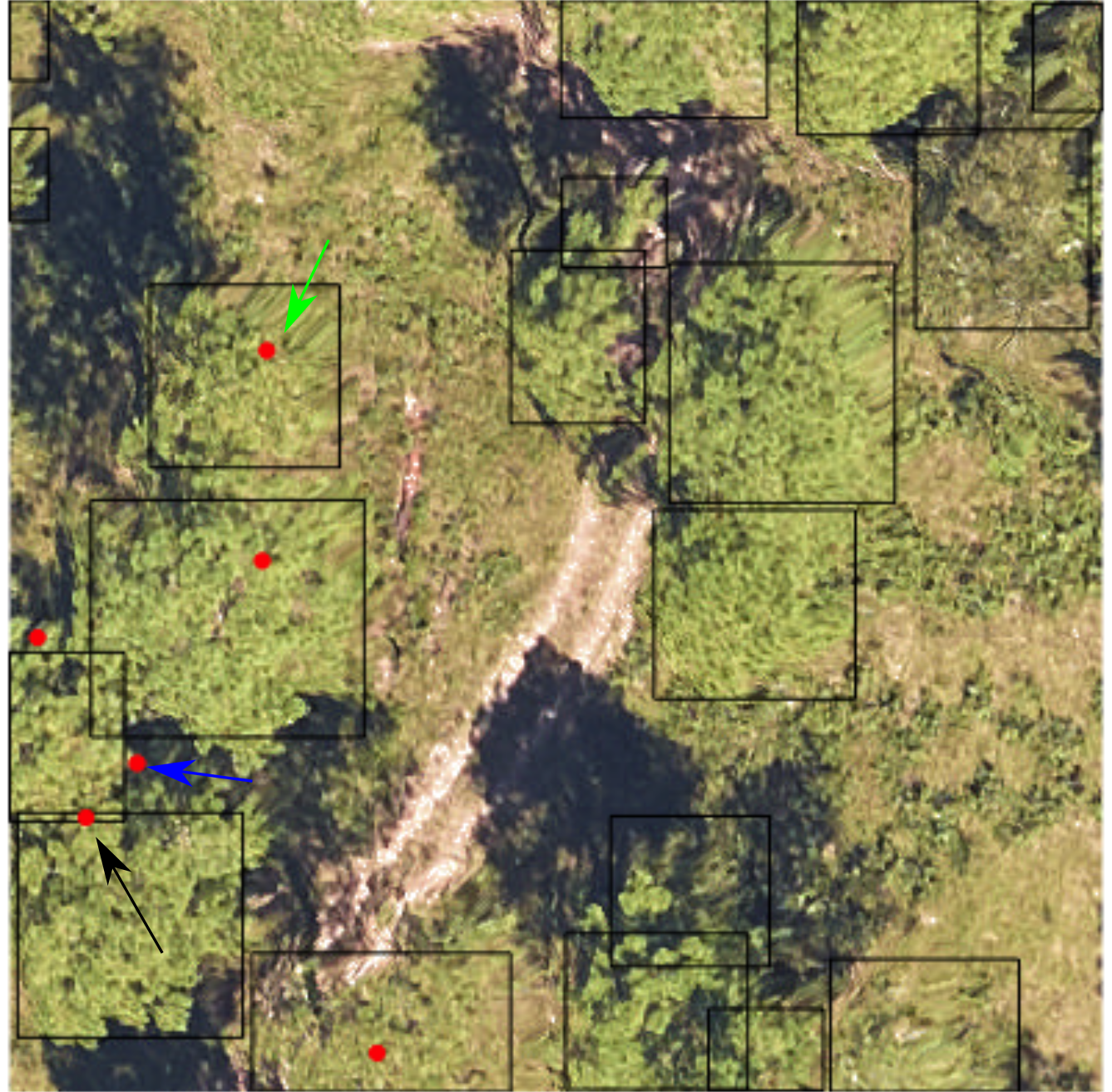

Figure S5. Example stem recall for evaluation plot JERC_048 from Jones Ecological Resource Center, Georgia. Predicted tree bounding boxes in black. Filtered NEON field stems (see above for filtering criteria) in red.

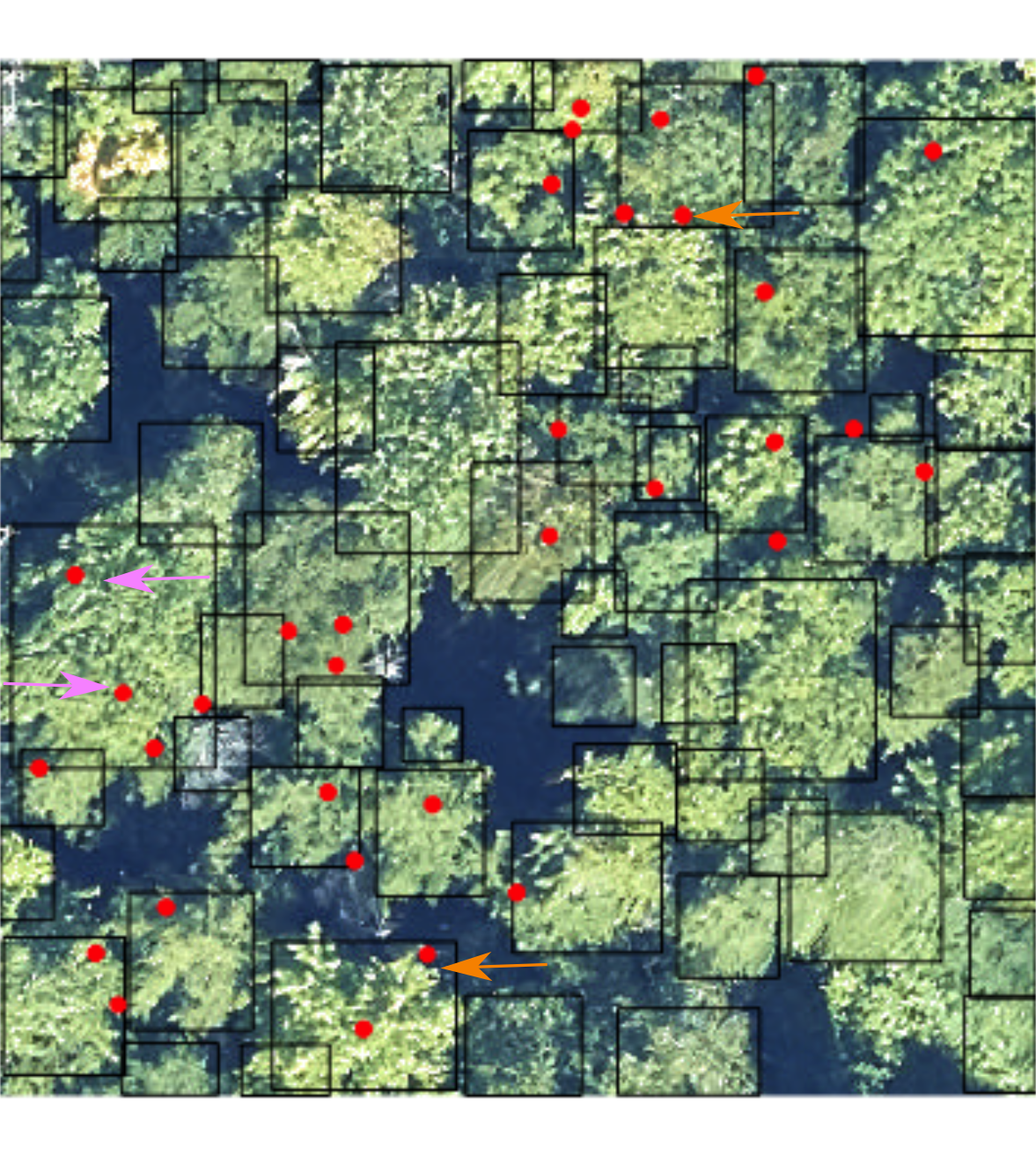

S6. Example from NEON plot BART_050 from Bartlett Forest, New Hampshire. In dense forests, multiple field stems can fall within a single predicted bounding box. This is due to both under-segmentation of visible crowns (pink arrows), as well as potentially incomplete filtering of field stems to identify overstory trees visible in the image (orange arrows).

Figure S5 and Figure S6 highlight some of the challenges of matching remote sensing crown predictions and field collected stem data. In Figure S5, the green arrow shows the simplest scenario, a single prediction unambiguously overlaps a single collected stem. The black arrow shows a moderately challenging scenario in which a visually unambiguous crown only barely matches the field collected stem. This could occur due 1) the stem growing at an angle leading to a spatial mismatch between crown and stem, 2) spatial error in measuring the crown location, 3) spatial error in the georeferencing of the RGB image. The blue arrow shows a stem point that does not overlap with a prediction crown. We have written the stem recall evaluation to be conservative, allowing no tolerance for points outside of the prediction box. Therefore, the stem recall for this image was 4/6 = 66.66%, which we believe is a conservative representation of the performance of the algorithm given the uncertainty in the field data and matching process.

### S2. Qualitative Assessment of Broad Scale Predictions

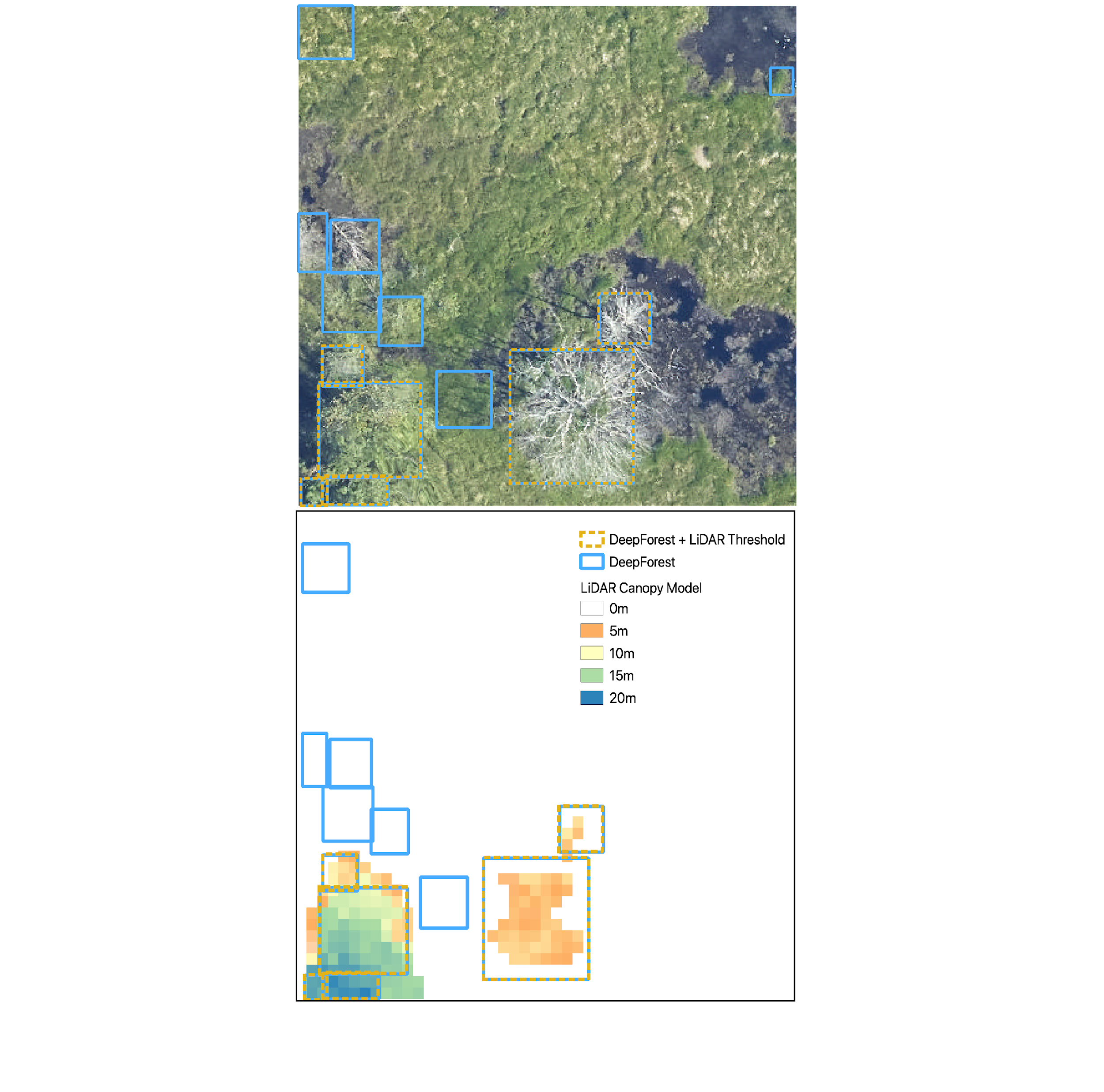

Figure 4. Illustration of the LiDAR threshold for minimum predicted tree height using NEON plot ID OSBS_023 from Ordway Swisher Biological Station, Florida. Using the DeepForest algorithm (Weinstein et al., 2020a), in blue, several boxes were removed based on no LiDAR returns above 3m (dotted orange). This step was key in open-ground areas in which the algorithm can confuse short vegetation with standing trees.

With over 7,000 1km2 tiles, it is not possible to do a systematic check of predicted crowns. Our aim is to provide users with a broad scale description of areas of concern. There are two main types of failure modes, data quality and algorithm quality. Data quality errors can occur in either the RGB data or LiDAR products. RGB errors include incomplete site coverage, image artifacts due to georectification, errors in orthomosaic stitching and lighting changes among data collection events (Figure S6).

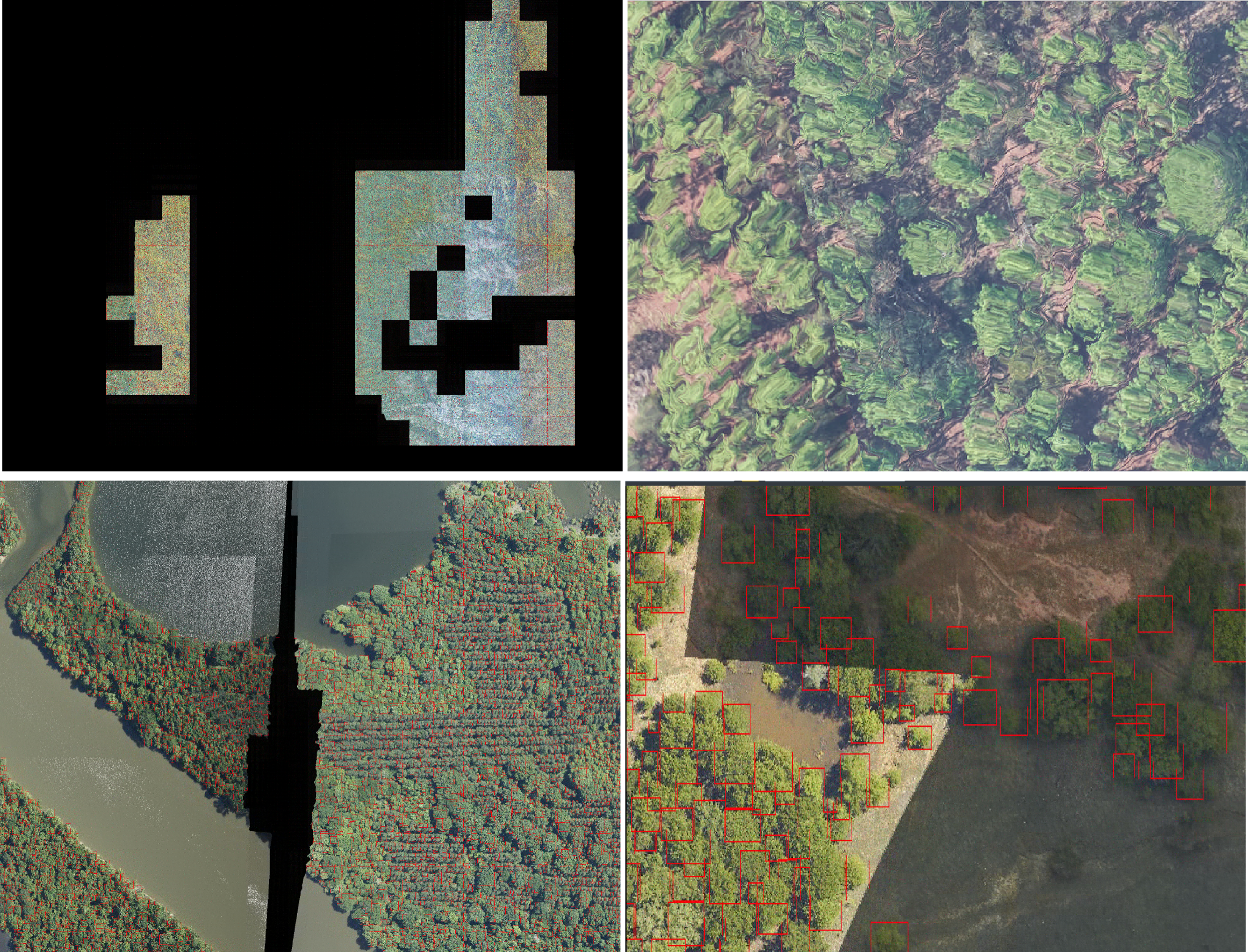
Figure S6. Errors in data quality leading to inaccurate predictions or inappropriate use cases. Top left) Patchy coverage at the site level can prevent broad scale analysis with large gaps in RGB data availability (NEON site GRSM). Top right) RGB artifacts at the edge of images stem from the challenge of georectification of the RGB and LiDAR tiles leading to swirling in the RGB pixels (NEON site OSBS). Bottom left) Seams in the flightlines in the RGB images leads to gaps in local predictions (NEON site DELA). Bottom right) Changes in illumination among data acquisition collections lead to stark changes in pixel values.

### S3. NEON Site Abbreviations

| **Site Name** | **siteID** | **Domain Number** | **State** | **Latitude** | **Longitude** |
| --- | --- | --- | --- | --- | --- |
| Abby Road | ABBY | D16 | WA | 45.76243 | -122.33033 |
| Bartlett Experimental Forest | BART | D01 | NH | 44.06388 | -71.28731 |
| Blandy Experimental Farm | BLAN | D02 | VA | 39.06026 | -78.07164 |
| Caribou-Poker Creeks Research Watershed | BONA | D19 | AK | 65.15401 | -147.50258 |
| LBJ National Grassland | CLBJ | D11 | TX | 33.40123 | -97.57 |
| Rio Cupeyes | CUPE | D04 | PR | 18.11352 | -66.98676 |
| Delta Junction | DEJU | D19 | AK | 63.88112 | -145.75136 |
| Dead Lake | DELA | D08 | AL | 32.54172 | -87.80389 |
| Disney Wilderness Preserve | DSNY | D03 | FL | 28.12504 | -81.4362 |
| Guanica Forest | GUAN | D04 | PR | 17.96955 | -66.8687 |
| Harvard Forest | HARV | D01 | MA | 42.5369 | -72.17266 |
| Healy | HEAL | D19 | AK | 63.87569 | -149.21334 |
| Lower Hop Brook | HOPB | D01 | MA | 42.47179 | -72.32963 |
| Jones Ecological Research Center | JERC | D03 | GA | 31.19484 | -84.46861 |
| Jornada LTER | JORN | D14 | NM | 32.59068 | -106.84254 |
| Konza Prairie Biological Station | KONZ | D06 | KS | 39.10077 | -96.56309 |
| Lajas Experimental Station | LAJA | D04 | PR | 18.02125 | -67.0769 |
| Lenoir Landing | LENO | D08 | AL | 31.85388 | -88.16122 |
| Mountain Lake Biological Station | MLBS | D07 | VA | 37.37828 | -80.52484 |
| Moab | MOAB | D13 | UT | 38.24833 | -109.38827 |
| Niwot Ridge Mountain Research Station | NIWO | D13 | CO | 40.05425 | -105.58237 |
| Northern Great Plains Research Laboratory | NOGP | D09 | ND | 46.76972 | -100.91535 |
| Klemme Range Research Station | OAES | D11 | OK | 35.41059 | -99.05879 |
| Ordway-Swisher Biological Station | OSBS | D03 | FL | 29.68927 | -81.99343 |
| Red Butte Creek | REDB | D15 | UT | 40.78374 | -111.79765 |
| Rocky Mountain National Park, CASTNET | RMNP | D10 | CO | 40.27591 | -105.54592 |
| Smithsonian Conservation Biology Institute | SCBI | D02 | VA | 38.89292 | -78.1395 |
| Smithsonian Environmental Research Center | SERC | D02 | MD | 38.89008 | -76.56001 |
| San Joaquin Experimental Range | SJER | D17 | CA | 37.10878 | -119.73228 |
| Soaproot Saddle | SOAP | D17 | CA | 37.03337 | -119.26219 |
| Santa Rita Experimental Range | SRER | D14 | AZ | 31.91068 | -110.83549 |
| Talladega National Forest | TALL | D08 | AL | 32.95046 | -87.39327 |
| Lower Teakettle | TEAK | D17 | CA | 37.00583 | -119.00602 |
| West St Louis Creek | WLOU | D13 | CO | 39.89137 | -105.9154 |
| Woodworth | WOOD | D09 | ND | 47.12823 | -99.24136 |
| Wind River Experimental Forest | WREF | D16 | WA | 45.82049 | -121.95191 |
| Yellowstone Northern Range (Frog Rock) | YELL | D12 | WY | 44.95348 | -110.53914 |
